# Self-assembly based post-translational protein oscillators

**DOI:** 10.1101/2020.02.12.946145

**Authors:** Ofer Kimchi, Carl P. Goodrich, Alexis Courbet, Agnese I. Curatolo, Nicholas B. Woodall, David Baker, Michael P. Brenner

## Abstract

Recent advances in synthetic post-translational protein circuits are significantly impacting the landscape of biomimicry engineering. However, designing sustained dynamic phenomena in these circuits remains an outstanding challenge. Inspired by the KaiABC system regulating the circadian clock in cyanobacteria, we develop two experimentally realizable post-translational oscillators. The oscillators rely on a small number of components interacting only through reversible binding and phosphorylation/dephosphorylation reactions.

## INTRODUCTION

Protein oscillators play a major regulatory role in organisms ranging from prokaryotes to humans. In most biological cases, the oscillation is realized through transcription/translation cycles. Few examples of purely post-translational oscillators have been found in biology [1, 2]. At the same time, post-translational protein circuits are increasingly sought after for synthetic applications, since they have the potential to exhibit faster response to environment changes, allow for more direct control over the circuit behavior, be directly coupled to a functional output, and can be employed in contexts that don’t include the vast genetic apparatus [3–5]. While significant recent work has enabled the design of post-translational protein-based logic gates [4, 5], engineering dynamic phenomena such as oscillations in a post-translational context remains an outstanding challenge [6, 7].

The best-studied example of biological post-translational protein oscillators is the KaiABC system in cyanobacteria [8]. By placing only the proteins KaiA, KaiB, and KaiC in a test tube, along with abundant ATP, the KaiC proteins collectively get sequentially phosphorylated and dephosphorylated, forming an oscillatory cycle [9, 10]. While the KaiC proteins generally exist in a hexameric state, monomers are shuffled among the hexamers during only a certain phase of the oscillatory cycle [11]. The KaiABC system demonstrates that protein oscillators need not use transcription/translation cycles or large numbers of components to achieve oscillatory behavior.

Motivated by the KaiABC system, we set out to design a protein-based oscillator that could be re-constituted *in vitro* using only a small number of components at relatively high copy numbers, so that any resulting oscillations are not stochastic. In order to facilitate the future translation of this theoretical study to an experimental system, we base the architecture of our system on biochemical constraints and on a design space navigable through computational protein design. We constrain the kinetic reaction network to only include three protein species, and only allow reversible binding and phosphorylation-dephosphorylation enzyme reactions.

The simplest such system, a protein with one phosphorylation site being modified by a kinase and a phosphatase, cannot yield oscillations regardless of parameter choices [7]. When two phosphorylation sites are included, oscillations are possible only under the assumption that each of the four possible phosphorylation states has significantly different rates of subsequent phosphorylations and dephosphorylations [7]. While biology seems to have designed a system in KaiABC capable of undergoing the many conformational changes necessary to implement this form of oscillations [9], the design of even two (let alone several) protein structures from the same sequence remains a significant challenge for the field of computational protein design [12].

These challenges are not unique to molecules with two phosphorylation sites. For example, since oscillations for molecules with two phosphorylation sites are effected by enzyme sequestration [7], we consider a molecule containing a single phosphorylation site alongside a kinase- or phosphatase-sequestering domain (or a binding domain for an external compound that itself contains an enzyme-binding domain). Such systems are capable of producing oscillations–but only if phosphorylation and binding accompany a significant conformational change in the molecule that modifies the rate constants of subsequent reactions. Even assuming such a conformational change were designed, we have found no evidence of sustained oscillations in such systems within the parameter regimes of typical binding/unbinding rate constants and typical kinase and phosphatase activity (i.e. the catalytic rate and Michaelis constants *k*_cat_ and *K*_*M*_, discussed further below). See Fig. S1 for further discussion.

Systems which focus on modifications to the enzymes themselves are therefore more likely candidates for the production of experimentally realizable oscillations. Biology has found several ways to tie phosphorylation to enzymatic activity. The most straightforward conceptually, having the activity of an enzyme dependent on its phosphorylation state [13] remains a challenge to implement in the context of computational protein design [14]. However, the field has achieved remarkable success in the design of protein-protein interactions [15] which can be modified by phosphorylation [16, 17]. Bootstrapping off of this success, we consider proteins which self-assemble into multimeric functional enzymes, motivated in part by the success of using split proteases to implement post-translational protein-based logic gates [4, 5]. In our design, when the proteins’ binding interfaces are phosphorylated, their self-assembly is impededed, reducing the concentration of functional enzymes available in the system.

Here, we describe two design schema for such self-assembly based post-translational protein oscillators. Our designed oscillators include only three protein species, and only allow reversible binding and phosphorylation-dephosphorylation enzyme reactions. We first consider self-assembly into closed symmetric homomultimers of specified size, and then discuss a similar system of one-dimensional unbounded assembly (fibers). These two system are described schematically in Fig. 1a and b respectively.

**FIG. 1.**
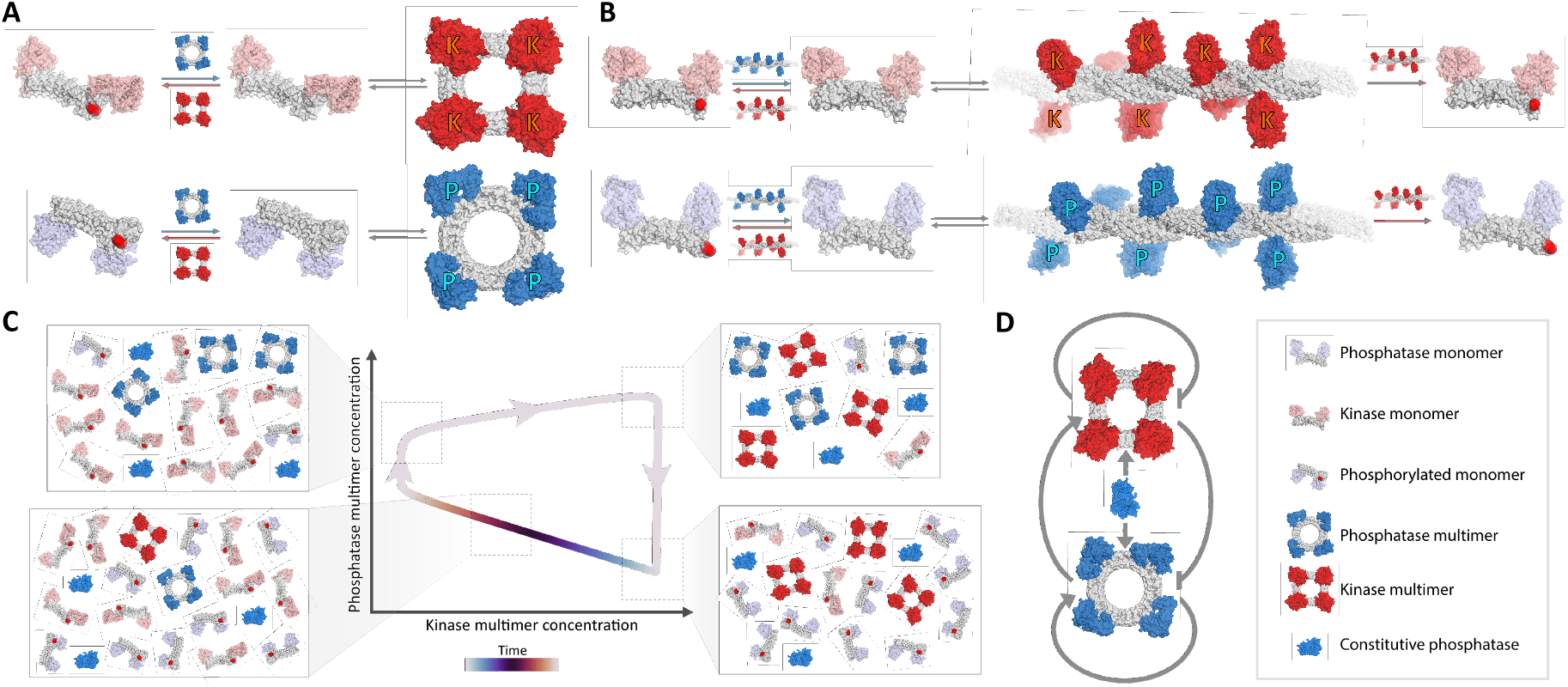
Model and oscillations overview. **A: Bounded self-assembly** (Eqn. 1). Monomers contain two halves of a split enzyme: either kinase (red; top) or phosphatase (blue; bottom). Monomers can self-assemble into multimers of specified size (here, tetramers are pictured, corresponding to *n* = *m* = 4). Kinase (phosphatase) multimers can (de)phosphorylate the monomers. A constitutive phosphatase is also able to dephosphorylate the monomers (not pictured). Phosphorylated monomers cannot participate in the self-assembly. **B: Unbounded self-assembly** (Eqn. 9). In this model, self-assembly of the kinase and phosphatase monomers is into unbounded fibers of arbitrary length. In addition, we assume the final monomer of each fiber can get phosphorylated by a kinase multimer, at which point it can no longer rejoin the fiber until it is dephosphorylated. **C: Oscillation schematic**. A sample oscillation of the simplified bounded self-assembly model (Eqn. 1) using arbitrarily chosen parameters satisfying experimental constraints. The system starts with self-assembled kinases and phosphatases (top right). The phosphatases disassemble much faster than the kinases, and get phosphorylated by the latter (bottom right). The kinases slowly disassemble, enabling the gradual dephosphorylation and self-assembly of the phosphatase monomers (bottom left). Once kinase levels fall below a critical threshold, the assembled phosphatases are able to rapidly promote their own self-assembly through dephosphorylation faster than the kinases can disrupt it (top left). Kinase monomers are then able to self-assemble and return the system to its initial state (top right). **D: Oscillator topology**. By phophorylating monomers, kinase multimers (red; top) inhibit their own and phosphatase multimer (blue; bottom) self assembly. Similarly, phophatase multimers counteracts this inhibition, as does constitutive phosphatase (center).

## RESULTS

### Self-assembly based protein oscillators are able to function within experimental constraints

The main components of our oscillators are two proteins, which we call *κ* and *ρ*. Each individual protein of type *κ*(*ρ*) has two complementary parts of a split kinase(phosphatase) and a phosphorylation site. When the respective sites are dephosphorylated, copies of protein *κ*(*ρ*) can self-assemble into a functional kinase(phosphatase), which we call *K*(*P*). Thus, self-assembled kinases inhibit the self-assembly of new proteins while self-assembled phosphatases counteract the inhibition (Fig. 1c). The resulting circuit topology (Fig. 1d) is analogous to that used in the dual-feedback genetic oscillator [18]. We treat the self-assembled enzyme as only one functional protein because the copies of the enzyme are all colocalized. In addition to the proteins *κ* and *ρ*, we include a constitutive phosphatase 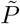; without it, a stable fixed point where all proteins are phosphorylated can preclude oscillations.

Because we are motivated by experimental feasibility, we consider only physically realizable parameters for our models. Binding rates *k*_*b*_ are typically in the range 10^−2^ − 10^0^ *μ*M^−1^*s*^−1^ [19] with dissociation constants *k*_*d*_ typically in the 10^−3^ − 10^3^ *μ*M range [20]. Both of these quantities can be tuned based on the geometry, energy, and symmetry of the binding interface between the proteins, which we assume here to be designed *de novo*. Less straightforward to design are the Michaelis constants and catalytic rates of the kinase and phosphatase, especially since these depend strongly not only on the enzyme but on the substrate. Mutational screens can be used to adjust the parameters but predicting the effect of a mutation on *k*_cat_ or *K*_*M*_ is highly nontrivial [21]. We were unable to find studies measuring kinase and phosphatase rates on the same substrate. Instead, as a standard to demonstrate physical realizability, we consider the parameters for sample Ser/Thr enzymes: wildtype *λ*-PPase (phosphatase) acting on pNPP (*k*_cat_ = 2.0 × 10^3^ *s*^−1^; *K*_*M*_ = 1.0 × 10^4^*μ*M) and wildtype MST4 (kinase) acting on the short peptide chain NKGYNTLRRKK (*k*_cat_ = 3.1 *s*^−1^; *K*_*M*_ = 14*μ*M) [21, 22]. We assume throughout that the constitutive phosphatase 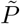 behaves identically to the self-assembled *P*.

To determine if the two systems are capable of producing oscillations within experimental constraints, we numerically integrated their respective kinetic equations within the parameter ranges outlined above. Our results, presented in Fig. 2, demonstrate a significant portion of parameter space in each system capable of admitting sustained oscillations. To our knowledge, these systems represent the first synthetic frameworks for experimentally realizable post-translational protein oscillators.

**FIG. 2.**
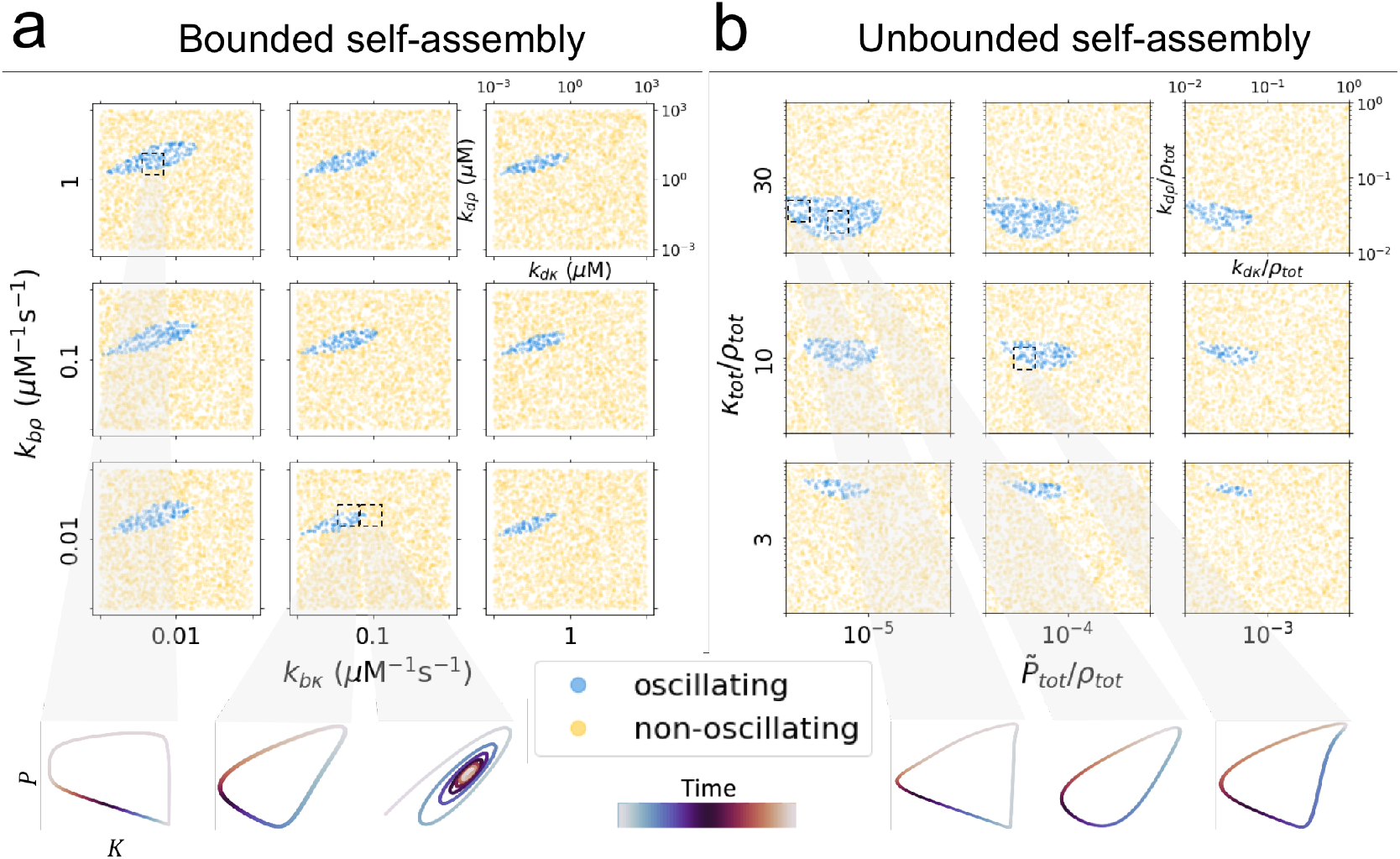
a: Bounded two-state self-assembly oscillations. Numerical integration of Eqn. S1 (Fig. 1A) displays parameter regimes leading to oscillations within experimental constraints. Each subplot shows the location of oscillating parameter sets as a function of *k*_*dκ*_ and *k*_*dρ*_ for given *k*_*bκ*_ and *k*_*bρ*_; the latter two are varied for each subplot. Aside from experimental constraints (see main text for discussion) we set *n* = *m* = 2, *κ*_tot_ = *ρ*_tot_ = 10*μ*M, and 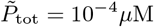. Blue points denote parameter sets leading to sustained oscillations; yellow points to steady state. Below the plot we show a few example trajectories in *K* – *P* phase space. We show that even minor variations in parameters can yield qualitatively different dynamics. In these trajectories, the values of the concentrations and time are omitted for clarity, but can be found in Fig. S2. **b: Unbounded incremental self-assembly oscillations**. Numerical integration of Eqn. 9 (Fig. 1B) displays parameter regimes leading to oscillations within experimental constraints. Eqn. 9 was used in place of the full system of equations (Eqn. S3) because of the infinite-dimensionality of the latter. *ρ*_tot_ sets the concentration-scale.

### Bounded self-assembly can yield oscillations whose behavior is well-predicted by simple limits

To find how the system parameters determine the characteristics of the oscillations, we seek simplified analytically tractable formulae that clarify the fundamentals of the oscillations. To this end, we consider concentrations which are low compared to the Michaelis constants, such that the concentration of enzymatic intermediates can be neglected. This approximation, like others we will consider, is not obeyed by all oscillating solutions found numerically (Fig. 2) but is nonetheless useful in clarifying the fundamentals of a large swath of the oscillations. We find that, in contrast to well-known examples from other systems which rely on enzyme sequestration to achieve oscillations [6, 7], neglecting enzyme sequestration does not preclude oscillations for our systems. In order to reduce our systems further to only two differential equations, we assume a separation of timescales between the self-assembly and the enzymatic activity.

For the first system we’ll describe of all-or-nothing bounded self-assembly (Fig. 1a), we assume that phosphorylation/dephosphorylation reactions equilibrate much faster than self-assembly. After accounting for conservation laws and the approximations described, we arrive at the following two-dimensional system of equations:

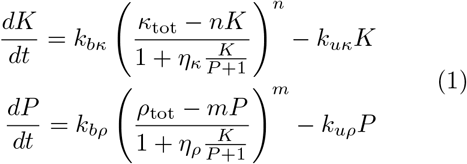

where we’ve normalized all concentrations (including binding rates) by dividing by 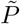. We briefly define the parameters: *n*(*m*) is the number of monomers in a multimer of *κ*(*ρ*); *k*_*bκ*_ is the binding rate for *κ* into its multimeric state, *k*_*uκ*_ is the respective unbinding rate, and *k*_*dκ*_ the inverse ratio of the two; *η*_*Kκ*_ is the specificity constant *k*_cat_/*K*_*M*_ for the kinase *K* acting on *κ*, *η*_*Pκ*_ is the same for the phosphatase *P*, and *η*_*κ*_ = *η*_*Kκ*_/*η*_*Pκ*_; *κ*_tot_ is the total concentration of monomeric *κ* added to the system, a conserved quantity. Similar quantities are defined for *ρ*.

In order to describe the oscillatory behavior of the system, we seek the eigenvalues of the Jacobian in the vicinity of a fixed point (*K*^⋆^, *P*^⋆^). Oscillations require coupling between the equations, motivating the approximations that in the oscillatory regime, *η*_*κ*_*K*^⋆^ ≫ *P*^⋆^ + 1 (and same for *η*_*ρ*_), *κ*_tot_ ≫ *nK*^⋆^, and *ρ*_tot_ ≫ *mP*^⋆^. The fixed point in these limits is given by

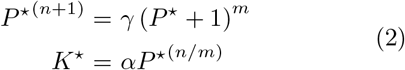

where *γ* and *α* are unitless constants defined by

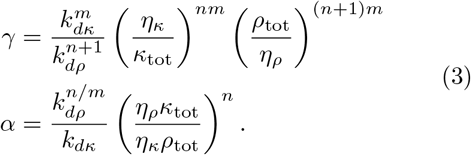

We constrain ourselves to *m* ≤ *n*+1 so that within our approximations there is no more than one physical fixed point in the system as long as 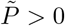, simplifying our analysis. (When no solutions to Eqn. 2 exist, our assumptions leading to it break down).

Sustained oscillations in the system typically correspond to complex eigenvalues of the Jacobian with positive real parts. However, following the Poincaré-Bendixson Theorem, as long as our system has a single fixed point, instability of the fixed point must imply oscillations even if they are beyond the linear regime. Translated into constraints on *P^⋆^*, instability of the fixed point gives a lower bound:

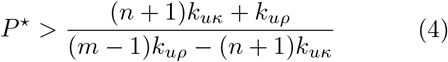

The upper bound on *P*^⋆^ is given by self-consistency with the previous assumptions we made (those leading to Eqn. 2).

Because Equation 2 can’t be solved for *P*^⋆^ for general *n, m*, we consider the approximation that *P*^⋆^ = *γ*^1/(*n*+1−*m*)^ ≫ 1, equivalent within the constraint *m* < *n* + 1 to *γ* ≫ 1. This approximation is most accurate for small values of *m*, since fewer terms are neglected. The approximation is motivated by the intuition that oscillations require *P* to be non-negligible compared to 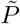; indeed, an opposite self-consistent solution, in which *P*^⋆^ ≪ 1, is incompatible with oscillations.

In this limit, oscillations can be found when

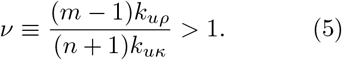

We verify that the approximate formula given by Equation 5 is valid in describing Eqn. 1 by comparing it to oscillations found by random parameter searches in panel a of Fig. 3. We numerically integrate the unitful version of Eqn. 1 with random parameters chosen to satisfy the experimental constraints described previously (including setting *η*_*κ*_ = *η*_*ρ*_) and with *n* = *m* = 2. We constrain con-centrations *κ*_tot_ and *ρ*_tot_ to be within 10^−3^ − 10^2^*μ*M, while we set the bounds of 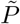 to 10^−8^ − 10^−2^*μ*M. For each parameter set, we numerically estimate the fixed point using Python’s scipy.optimize.root function. We only show parameter sets estimated to agree with the approximations described before Eqn. 2 (with > 5× substituted for ≫). We found no oscillations in ~ 2.5 × 10^4^ parameter sets which violate any of these assumptions (e.g. for which *η*_*K*_*K*^⋆^ < *P*^⋆^ + 1). Each blue (yellow) point in the figure corresponds to a single parameter set found to produce (not produce) oscillations starting from initial conditions of (*K, P*) = (0, 0). Oscillations are almost exclusively found in the quadrant *γ* > 1, *ν* > 1. Values of *γ* slightly less than unity are also found to produce oscillations, as shown in the figure.

**FIG. 3.**
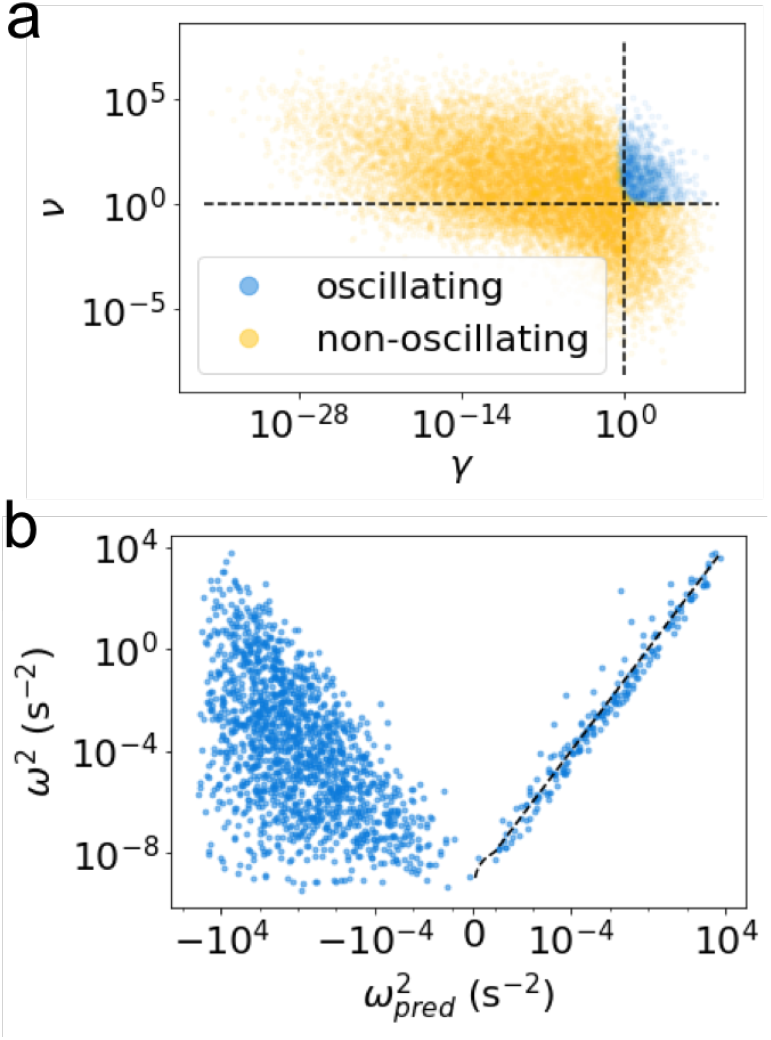
a: Onset of oscillations. Numerical integration demonstrates consistency with Eqn. 5 for the appearance of oscillations in the appropriate limits. Each point represents a random set of parameters, sampled within the experimentally realizable limits as described in the main text, with *n* = *m* = 2. Oscillating (blue) and non-oscillating (yellow) parameter sets can be well-separated by unitless combinations of parameters. Dashed lines show where the unitless parameters on the axes equal unity. **b: Oscillation frequency.** Numerical integration demonstrates Eqn. 6 correctly predicts the frequency of oscillations in the linear regime around the fixed point (dashed line represents 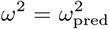), but is not predictive outside this regime.

In contrast to intuition from other systems which signifies that higher order nonlinearities increase the parameter range producing oscillations [6], here we found that more non-linear self-assembly (i.e. higher values of *n* and *m*) makes oscillations less frequent.

We next seek to predict how system parameters tune the frequency of resulting oscillations when they appear. Within the linear regime around the fixed point, in the limit of Eqn. 5, the frequency of oscillations *ω* is predicted to be

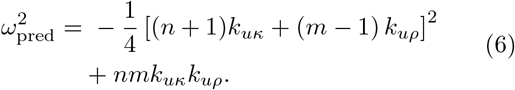

We compare Eqn. 6 to the true squared frequency for those parameter sets found to produce oscillations through numerical integration in panel b of Fig. 3. We make no constraints on the fixed points of the parameter sets considered here. The x-axis shows the predicted squared frequency while the y-axis shows the true squared frequency. For 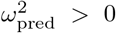, the two formulae agree (dashed line). For 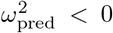, 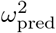 is no longer predictive since the oscillations cannot be understood through linear stability analysis of the fixed point. Oscillations found within experimental constraints for Eqn. 1 have periods ranging from fractions of a second to > 1 day (Fig. S3). For oscillations found for the full system of equations plotted in Fig. 2a, we find periods within a slightly more constrained range than for the simplified system but still spanning orders of magnitude, between ~ 1 minute and > 1 day (not shown).

### Unbounded self-assembly can yield oscillations within experimental constraints in an opposite limit

We next consider the second system in which individual species *κ* and *ρ* can self-assemble incrementally into one-dimensional unbounded fibers (Fig. 1b). While in the previous system, no phosphorylation sites are accessible in the multimeric state, in this system, one is (corresponding to the final protein in the fiber). An *n*-mer of species *X*, *X*_*n*_, can be created either from binding two smaller molecules *X*_*m*_ and *X*_*n*−*m*_, or from the spontaneous breaking of a bond of a larger molecule. A molecule of *X*_*n*_ can be destroyed either by binding to any other molecule or breaking any of its *n* − 1 bonds. The equations for self-assembly of species *X* are therefore given by

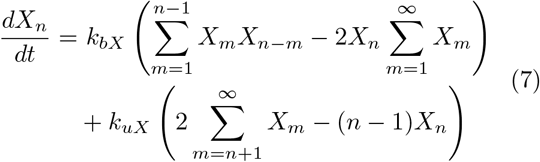

As in the first system, each protein of type *κ*(*ρ*) includes a split kinase(phosphatase). We assume when a multimer is phosphorylated its final monomer dissociates from the fiber and cannot reassociate in its phosphorylated state. A less stringent assumption, that the phosphorylated monomer doesn’t dissociate automatically but merely prevents new monomers from binding to that end of the molecule, appears to be incompatible with oscillations, at least in both separation-of-timescales limits.

While the limit considered for the first system, of fast phosphorylation/dephosphorylation compared to self-assembly, is no longer applicable for this system, we find oscillations in the opposite limit, of fast self-assembly compared to enzymatic activity. At steady-state, *X*_*n*_ is given by:

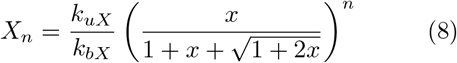

where *x* = 2*k*_*bX*_ *X*_*tot*_/*k*_*uX*_ = 2*X*_*tot*_/*k*_*dX*_. The same steady state is reached even if self-assembly involves binding and unbinding only a single monomer at a time.

The concentrations of phosphorylated monomers as a function of time are given by *κ*^*p*^ and *ρ*^*p*^. The total amount of kinase present is given by 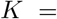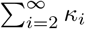, and similarly for phosphatase. Since only the phosphorylation site of the final monomer in a multimer is exposed, the total number of available phosphorylation sites in the *κ* species is given by 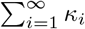 (and similarly for *ρ*). After neglecting intermediates as previously, the system can be described by two differential equations for *k* = 2(*κ*_tot_−*κ*^*p*^)/*k*_*dκ*_ and *p* = 2(*ρ*_tot_ − *ρ*^*p*^)/*k*_*dρ*_ (as in Eqn. 1 we normalize concentrations by 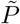):

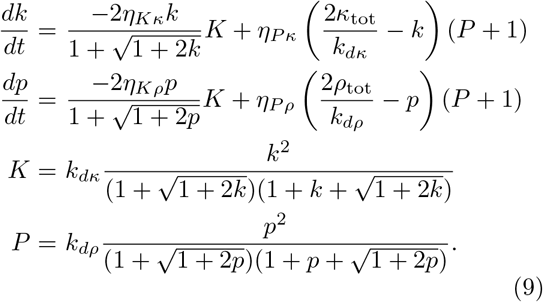

Unlike in the previous system for which the frequency and onset of oscillations can be determined by simple formulae by taking a limit of the two-dimensional system, no such limits give similarly straightforward results for Eqn. 9. Instead, we analyze Eqn. 9 through random parameter searches (Fig. 2b). We chose to analyze this two-dimensional system directly rather than the full equations (Eqn. S3) because of the infinite-dimensionality of the latter. As shown in the figure, we find that increased values of *κ*_tot_/*ρ*_tot_ and decreased values of 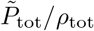 lead to more robust oscillations in this system. The periods of oscillations found ranged from < 1 minute to > 1 day (Fig. S3).

## DISCUSSION

In summary, we have presented two post-translational protein-based oscillators motivated by the biological KaiABC system. Both systems we present rely on split kinase and phosphatase self-assembling to form functional enzymes, and on that self-assembly being inhibited by phosphorylation of the split monomers. The two systems differ mainly in the nature of the self-assembly as all-or-nothing into bounded structures of specified size or incremental into unbounded one-dimensional fibers.

Both systems are capable of producing oscillations within experimental constraints, using experimentally-determined wildtype values for kinase and phosphatase activity and for a range of designed self-assembly rates. We have shown that neither complex reactions nor large number of species are necessary to achieve oscillations: both networks we present use only three protein species interacting only through reversible binding and phosphorylation/dephosphorylation reactions, and the resulting oscillations can be understood as arising from a minimal system of two differential equations in both cases. Although the systems we described shared much in common, they only produced oscillations in limits which precluded oscillations in the other system: the first oscillates when self-assembly is much slower than enzymatic reactions; the second when it is much faster.

Our work paves the way towards the rational design and experimental realization of protein-based far-from-equilibrium dynamic systems. The models described here were designed to be feasible to synthesize experimentally, and are guiding an implementation in the test tube that is currently underway.

We thank Arvind Murugan for helpful discussions.

This work was supported by the Harvard Materials Research Science and Engineering Center Grant No. DMR-1420570 and ONR Grant No. N00014-17-1-3029 (to M.P.B). O.K. acknowledges funding from the DoD through NDSEG Fellowship 32 CFR 168a, the NSF-Simons Center for Mathematical and Statistical Analysis of Biology at Harvard award number 1764269, and the Harvard Quantitative Biology Initiative. A.C. is a recipient of the Human Frontiers Science Program Long Term Fellowship and a Washington Research Foundation Senior Fellow. M.P.B. is an investigator of the Simons Foundation.

The authors declare no competing interests.

## SUPPLEMENTARY MATERIAL

### Full kinetic equations

We denote the concentration of phosphorylated (monomeric) *κ* by *κ*^*p*^ (and similarly for *ρ*). The concentration of the enzyme-substrate complex comprised of *κ* and *K* bound is denoted *κ* · *K*. Binding, unbinding, and catalytic rate constants for the enzyme-substrate complexes are given by *k*_*bKκ*_, *k*_*uKκ*_, and *k*_*cKκ*_, respectively. We use similar conventions for all other enzyme-substrate complexes. The full equations for the first system are:

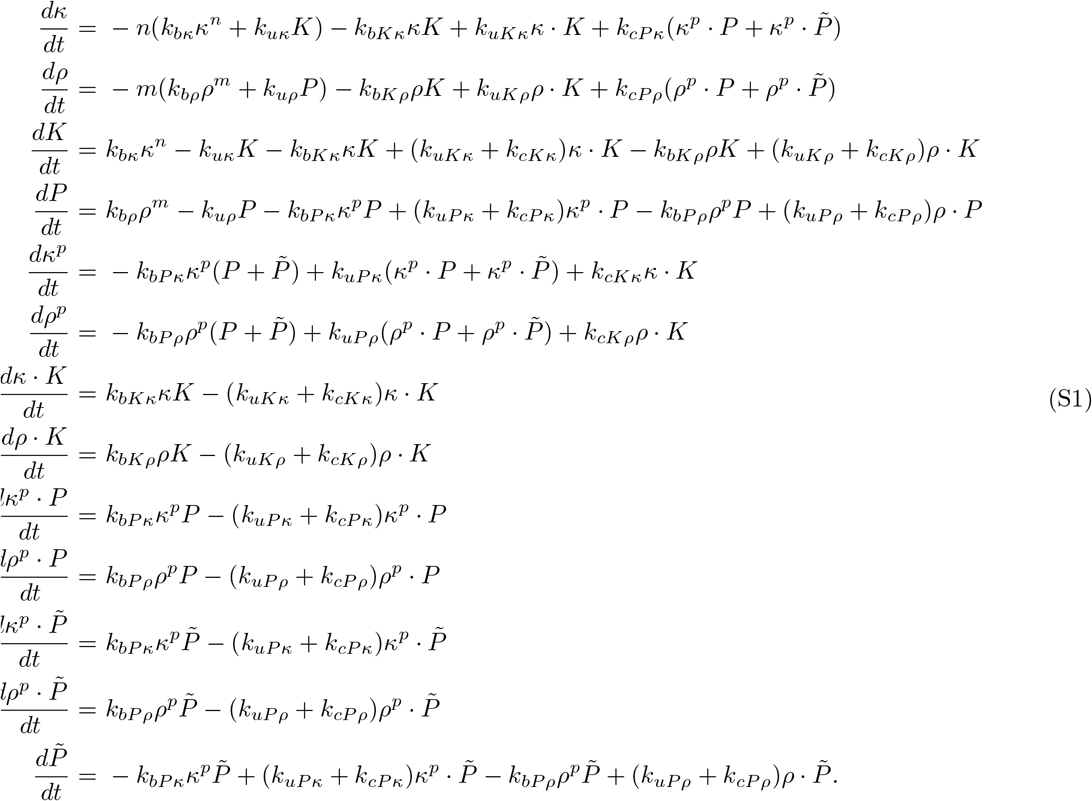

Making only the Michaelis-Menten approximation for enzymatic reactions and accounting for conservation laws, the equations can be reduced to the following four-dimensional system of equations:

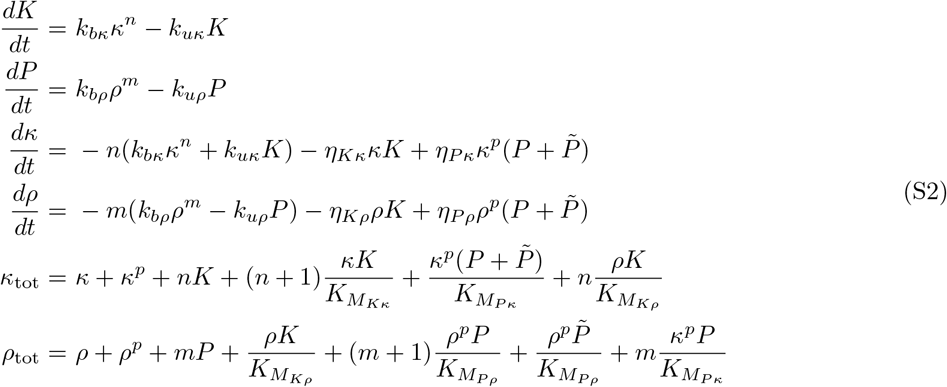

The full equations describing the second system are:

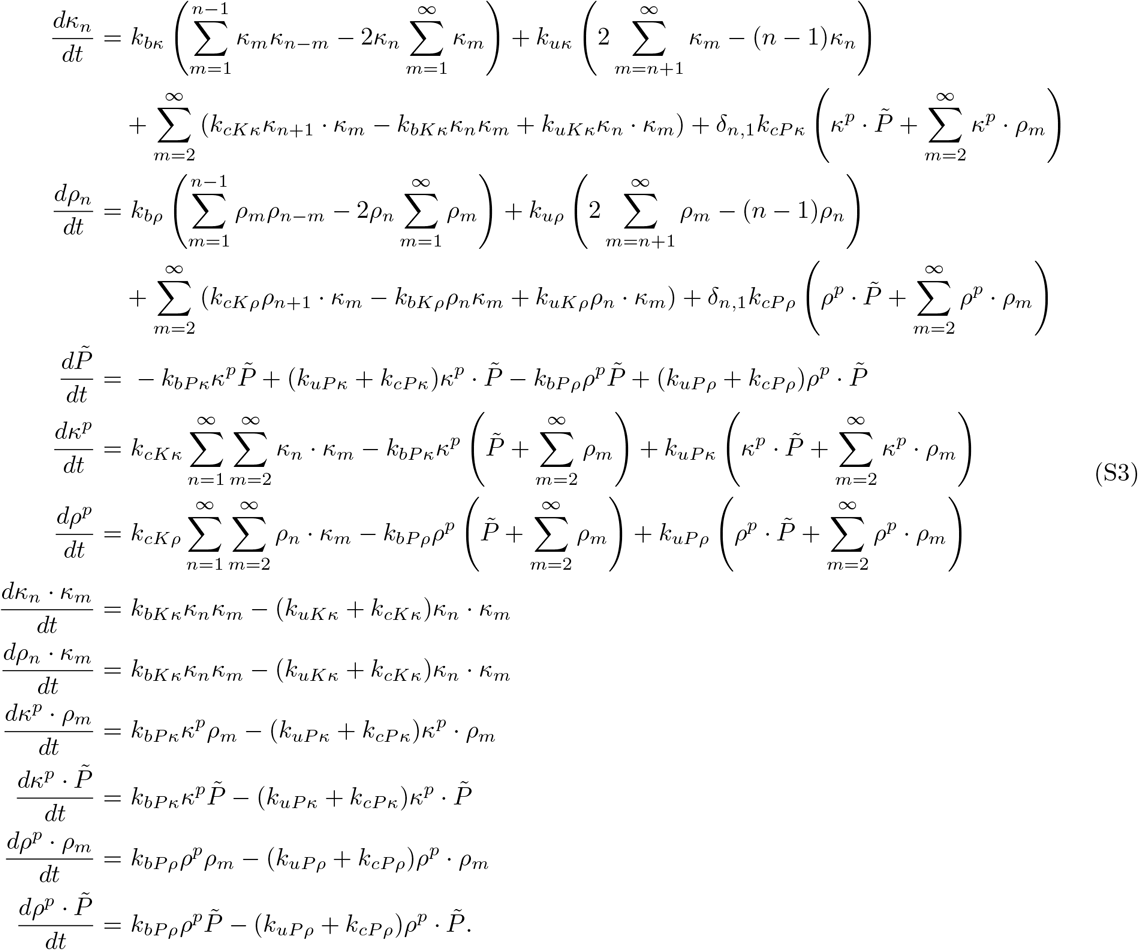

Within the Michaelis-Menten approximation and after accounting for conservation laws, these equations become:

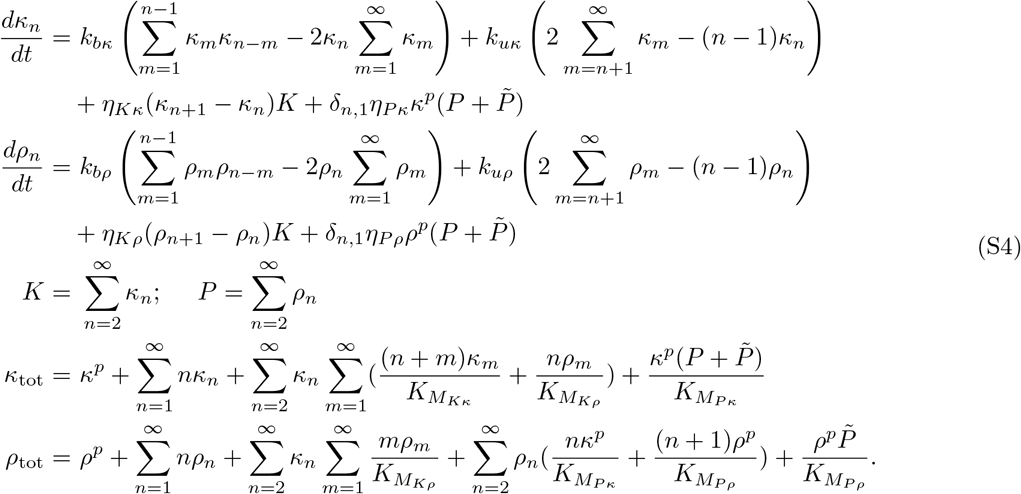

**FIG. S1.**
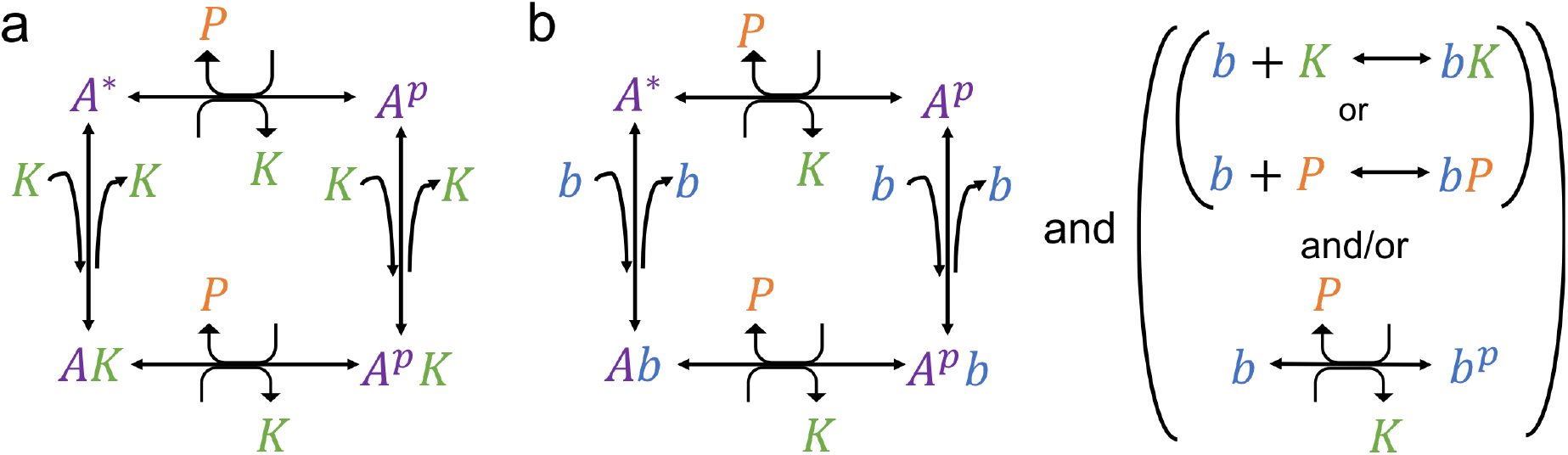
Reaction networks giving oscillations outside of experimentally realizable regime. See text for discussion.

In order to arrive at Eqn. 9, we assume a separation of timescales between the self-assembly and the enzymatic activity. In particular, we assume that self-assembly reactions equilibrate much faster than phosphorylation/dephosphorylation; the opposite limit does not seem to allow oscillations. We can then write the dynamics of the system only in terms of the phosphorylated monomers *κ*^*p*^ and *ρ*^*p*^.

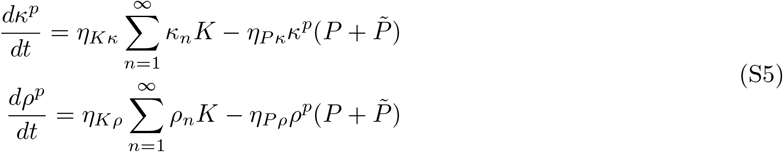

where *K* and *P* are as defined in Eqn. 9. Writing these equations in terms of *k* and *p* and normalizing concentrations by 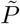, we arrive at Eqn. 9.

### Other oscillation schemes attempted

Before trying self-assembly based oscillations, we tried implementing oscillations based on phosphorylations or binding events accompanying a conformational change in the molecule. Such conformational changes can be difficult to design, but the recently-published LOCKR system [23, 24] demonstrates one way in which binding can accompany a conformational change. We considered a molecule *A* which can be phosphorylated or bind to another molecule. We assume that when it is bound or phosphorylated, the molecule undergoes a conformational change; in the language of the LOCKR system, it opens. We assume that the rate of binding of any molecule to the closed state *A*^⋆^ can be smaller than the analogous binding rate to the open state, but no other asymmetries between the rates of analogous reactions are allowed. We did not make simplifying assumptions such as the Michaelis-Menten approximation when considering these systems.

We found oscillations are possible if *A* can bind, and thus sequester, free kinases (Fig. S1a). Oscillations are also possible if *A* can bind a separate “key” peptide *b*, which itself either binds free kinase (*K*) or phosphatase (*P*) molecules. Finally, oscillations can also be found if *b*, either alongside or instead of binding kinase or phosphatase, can itself get phosphorylated. We assume phosphorylated *b* is inert, except in that it can interact with phosphatase to get dephosphorylated (Fig. S1b). However, we found no evidence of possible oscillations within the experimental limits considered in this paper, after trying for each network 2 × 10^6^ random parameter sets logarithmically distributed within the acceptable ranges.

### Numerical search for oscillations

In order to determine if a parameter set leads to oscillations, we numerically integrated the differential equations. For bounded self-assembly, we used initial conditions of (*K, P*) = (0, 0), and for unbounded, 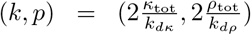. We integrated up to a time determined by the inverse of the minimum timescale in the system. For bounded self-assembly, we integrated up to a time 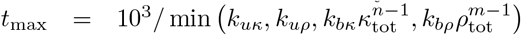, while for unbounded, we used *t*_max_ = 10^7^/ min (*η*_*Kκ*_*κ*_tot_, *η*_*Kρ*_*κ*_tot_, *η*_*Pκ*_*ρ*_tot_, *η*_*Pρ*_*ρ*_tot_). For the full system of equations for bounded self-assembly, we used initial conditions corresponding to fully unphosphorylated and unbound *κ*, *ρ*, and 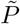, and integrated up to 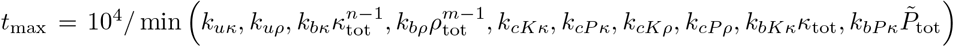. We set the enzyme dissociation constants *k*_*uKκ*_, *k*_*uPκ*_, *k*_*uKρ*_, *k*_*uPκ*_ equal to their respective catalytic rate constants, since the former are largely unspecified by constraints on binding rates and Michaelis constants. Our results are largely insensitive to this assumption. In all cases, the prefactors for *t*_max_ were determined by applying an order of magnitude larger prefactor and finding no new oscillating solutions.

To determine if the results of the numerical integration can be labeled as oscillations, we used a set of heuristics. We verified these heuristics by plotting solutions found by them to produce oscillations and finding no evidence of false positives. These heuristics considered the behavior of a single system component (e.g. *K* for bounded self-assembly). First, we determined whether the number of inflection points in the solution is greater than 10. Second, to weed out decaying oscillations, whether the smallest amount by which the component changed between inflection points and the amount it changed between an arbitrarily chosen set of inflection points (between the fifth and sixth) is within 2×. Also to weed out decaying oscillations, we measured the amount the component changed between a set of inflection points around the 3*t*_max_/4 mark–let’s call this amount *x*_3/4_–and between the penultimate and final inflection point, *x*_1_. We verified that |(*x*_3/4_ − *x*_1_)/*x*_1_| < 1, meaning that the relative change in oscillating height was no more than 100%. We also considered whether the solver required sampling points at a significant frequency (to weed out numerical oscillations): we used the criterion that the third-to-last sampled time point was within 5% of the second-to-last sampled time point. To further root out spurious numerical oscillations we measured the period of the oscillation in two ways–as the time between the third-to-last and last inflection point, and as between the fifth-to-last and third-to-last–and verified that they differed by no more than 1%. Finally, we examined the numerical solution by eye for all parameter sets found to produce oscillations, in order to verify that even if our heuristics produce false negatives (of which we have found almost no evidence) our results contain no false positives.

**FIG. S2.**
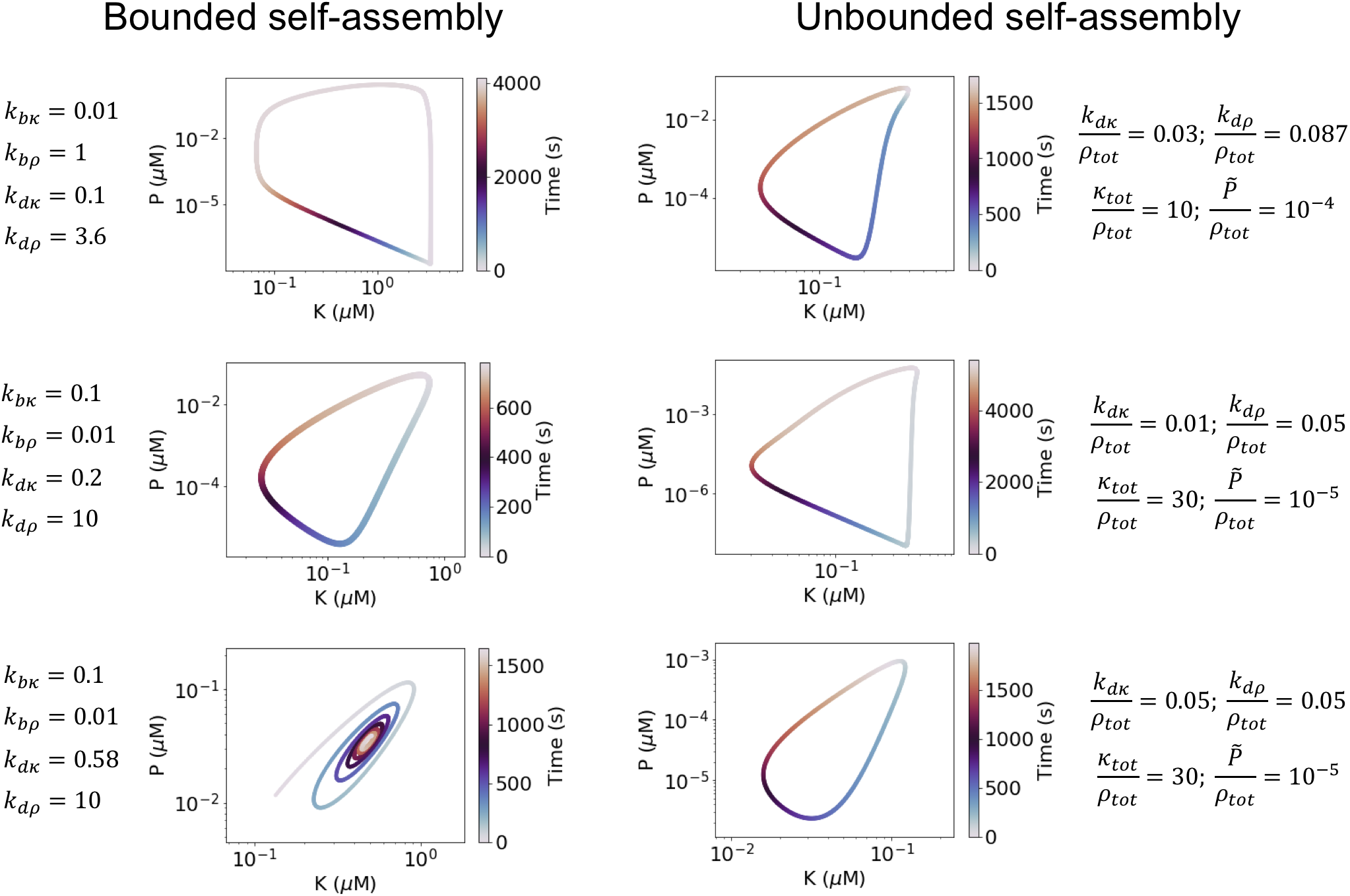
Example trajectories. Trajectories displayed in Fig. 2 are shown. We show the parameters used for each trajectory, the values of *K* and *P* along the trajectory, and time along the trajectory. For trajectories showing sustained oscillations (all but the lower left) one oscillation cycle is shown.

**FIG. S3.**
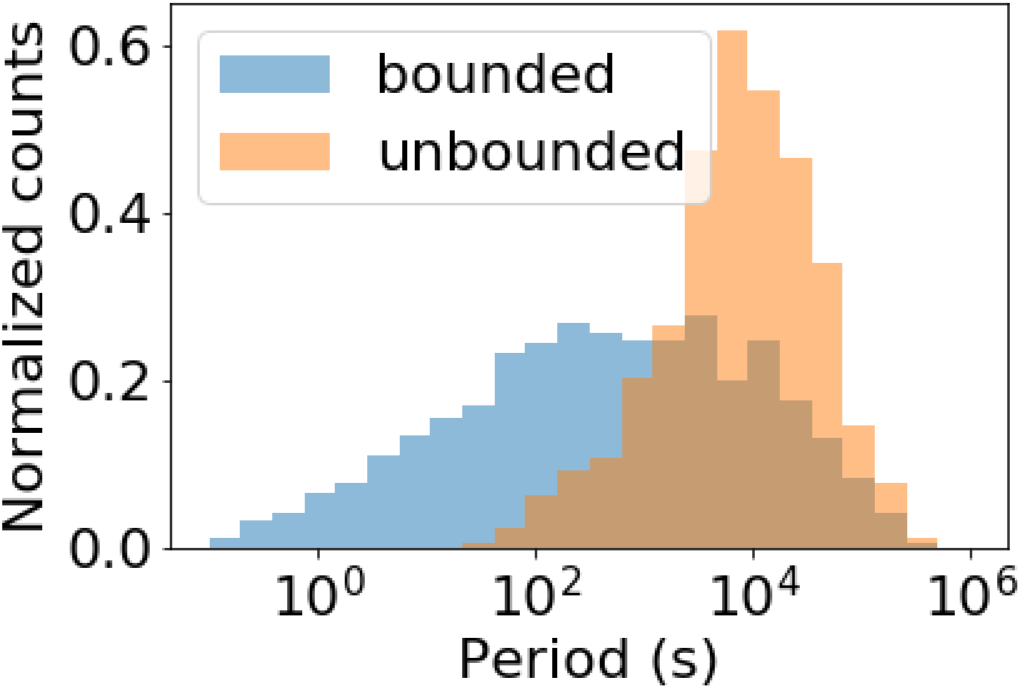
Oscillation periods. Random parameters logarithmically distributed within the experimental regime were sampled for Eqns. 1 (bounded self-assembly; blue) and 9 (unbounded self-assembly; orange). The periods of resulting oscillations are histogrammed logarithmically, showing a possible range of periods spanning orders of magnitude, from fractions of a second (minute) for bounded (unbounded) self-assembly, to > 1 day.

